# Bone Mineralization and Metabolism are Altered in a Rat Model of Brachial Plexus Birth Injury

**DOI:** 10.1101/2025.05.22.655603

**Authors:** Emily B. Fawcett, Jennifer R. Potts, Nikhil N. Dixit, Katherine R. Saul, Jacqueline H. Cole

## Abstract

Brachial plexus birth injury (BPBI) is a common nerve injury incurred during a difficult childbirth when the brachial plexus nerve bundle is excessively stretched, resulting in functional arm impairment in 30-40% of those affected. Injury can present in two different locations, modeled in rats as postganglionic and preganglionic neurectomies. Osseous deformities are present following both injury types. However, the underlying factors behind these deformities are not fully understood. While past studies have explored muscle structure and altered mechanical joint loading as factors, bone metabolism, muscle composition, and muscle-bone crosstalk have not been fully explored. Using postganglionic and preganglionic BPBI rat models and a disuse model, bone metabolism, muscle composition, and muscle-bone crosstalk were explored. Dynamic histomorphometry and similar methods were used to characterize humeral growth and humeral growth plate activity to understand bone metabolism, muscle fibrosis was analyzed to assess muscle composition, and FGF-2 quantification was performed to assess muscle-bone crosstalk. Postganglionic injury portrayed more changes in the humeral diaphyseal region than preganglionic and displayed reduced bone metabolism on the endosteal surface while preganglionic displayed reduced bone metabolism on the periosteal surface. However, only preganglionic showed significantly lower growth plate activity. In regards to fibrosis, both injury types showed fibrosis in the biceps but only preganglionic showed fibrosis in the subscapularis. The limb disuse model did not show fibrosis. Additionally, preganglionic had an increased production of FGF-2 signaling more so in the subscapularis. Overall, deformities from postganglionic injury may be from bone formation and bone resorption while deformities from preganglionic injury are likely from an overall reduction in bone growth that is not solely from limb disuse. The fibrosis and FGF-2 signaling alterations seen are not likely to be the direct cause of osseous deformity and the drivers behind the alterations are likely different between postganglionic and preganglionic injuries.

## Introduction

Brachial plexus birth injury (BPBI) occurs in 1-3 per 1000 births, making it one of the most common nerve injuries in children *(1)*. Injury occurs during an emergency birth when the head and shoulder are excessively stretched apart *(2)*, resulting in damage to the brachial plexus nerve bundle. In 30-40% of cases, lifelong arm impairment ensues *(3)*, causing muscle paralysis, osseous and postural deformity *(4)*, limb disuse *(5, 6)* and contractures *(5, 6)*. However, severity of shoulder consequences is dependent on the type of nerve injury *(7)*. A nerve rupture produces limb disuse and contractures *(8)* while a nerve avulsion produces limb disuse without contractures *(9)*. Regardless of injury type, treatment options are the same *(10)* and not consistently successful *(11)*, in part due to lack of knowledge surrounding shoulder deformities *(10)*.

Larger deformities, such as morphological bone changes and architectural muscle changes have been researched, but the reasoning for how and why the deformities occur have not been investigated. Using a murine model, nerve ruptures are represented by a postganglionic neurectomy *(6)* and nerve avulsion by a preganglionic neurectomy *(5)*. Both present with muscle alterations and osseous deformity, but the degree to which deficits are present varies *(5)*.

In regard to bone, postganglionic and preganglionic injuries show major differences *(12)*. Postganglionic results in morphological changes to the scapular glenoid region at a macrostructural level, including increased glenoid inclination angle *(6, 12-14)* and flattened curvature *(8, 12, 14)* that are not seen in preganglionic *(12, 14)*. However, preganglionic produces macrostructural changes to the humerus, such as a flattened *(8)* and smaller humeral head *(12)*, that are not seen in postganglionic. Additionally, postganglionic and preganglionic result in shorter humeri on the affected side *(6, 15, 16)*. At a microstructural level, postganglionic shows less robust trabecular bone in the humeral head *(16-19)* and glenoid fossa, but preganglionic shows even worse trabecular bone in both. While some of these deformities could be from disuse, there must be additional underlying factors.

In regard to muscle, postganglionic and preganglionic injuries produce smaller and shorter muscles *(5, 16, 17, 20-22)* with increased sarcomere length *(20, 22)*, but preganglionic shows more severe changes *(20)*. The composition of the muscle has been assessed only in the biceps, regardless of the fact that other muscles, such as the subscapularis muscle, show significant architectural changes. The fibrosis found in the biceps is similar between the two nerve injuries *(5)*, but the fibrosis in other muscles is unknown.

It has been theorized that the altered muscles following a postganglionic BPBI cause deformities through altered loading on the joint *(14, 23-26)*, producing growth into incorrect morphology, but other factors like bone metabolism and cellular crosstalk have not been investigated. Additionally, no studies specifically examine differences between the factors affecting postganglionic and preganglionic osseous deformity. The deformity drivers behind the postganglionic and preganglionic injuries could initiate changes in bone metabolism, muscle composition, or muscle-bone crosstalk (cell-signaling of growth factors). Comparing the two injury types will elucidate contributions from muscle contracture. Their comparison with a disuse model will then elucidate contributions of limb disuse.

Muscle-bone crosstalk occurs when muscle interacts with bone by secreting local growth factors which influence bone metabolism *(27)*. The interaction between the two can be affected via muscle composition. One study has examined muscle composition through the presence of fibrosis in the biceps and brachialis muscles *(5)*. However, the subscapularis muscle is also consistently involved in deformities *(20, 22)* and should be assessed. The study also looked at the preservation of ErbB signaling *(5)*. ErbB signaling is involved in myogenesis and muscle regeneration, both of which are essential to recovery after nerve injury. Signaling was shown to be preserved after a preganglionic injury but disrupted after a postganglionic injury *(5)*. However, this signaling is limited to within the muscle. Investigating a growth factor secreted by muscle such as FGF-2, which affects bone metabolism, and how it changes in regard to muscle composition, will give better insight to the muscle-bone interaction and how it changes with injury type.

The objective of this study is to investigate underlying bone metabolism, muscle composition, and muscle-bone crosstalk via FGF-2 after postganglionic and preganglionic injuries and the contributions from limb disuse. Examining these potential mechanisms will allow research into more effective treatment and therapy options which consider injury type differences. We hypothesize that alterations in bone growth and metabolism are primarily affected by muscle contracture and are associated with changes in both muscle composition and muscle-bone crosstalk. In other words, we hypothesize that postganglionic injury will have more severe alterations in bone metabolism, muscle composition, and muscle-bone crosstalk.

## Methods

### Study Design

All procedures were approved by the Institute of Animal Care. Forty-six Sprague Dawley rat pups (Harlan Laboratories, Indianapolis, Indiana) were split into three neurectomy groups: sham neurectomy (n=14), postganglionic neurectomy (n=16), and preganglionic neurectomy (n=16). An additional disarticulation group (n=12), which underwent amputation at the elbow, was used as a disuse model to compare to the nerve injury groups. All interventions occurred 3-5 days postnatal. In the postganglionic neurectomy group, a nerve rupture was mimicked by inducing a C5-C6 nerve root excision distal to the dorsal root ganglion *(8)*. Neurectomies were performed via incision through the pectoralis major (8). In the preganglionic neurectomy group, a nerve avulsion was mimicked by inducing a C5-C6 nerve root excision proximal to the dorsal root ganglion *(5)*. These neurectomies were performed via supraclavicular incision *(5)*. However, due to difficulty in visually confirming a full preganglionic neurectomy, some resulted in a partial nerve excision and were therefore excluded. This left 21 total for the preganglionic group. Criteria for incurring a full nerve excision included the phenotypic posture of an internally rotated shoulder and flexed wrist. The sham group underwent incision through the pectoralis major without any damage to the brachial plexus nerve bundle.

After intervention, animals were given one prophylactic dose of buprenorphine and the disarticulation group was given carprofen and all were weaned at postnatal day 21. Fluorochrome injections of calcein and alizarin were administered at 30 mg/kg body weight via IP injection at ten and three days before sacrifice respectively. At eight weeks post intervention, rats were sacrificed via CO2 inhalation and tissue of the affected and unaffected limbs were harvested. Immediately following dissection of tissue, the biceps short head, biceps long head, upper subscapularis, and lower subscapularis muscles were snap frozen in 2-methylbutance and stored at -80 degrees Celsius. Humeri and scapulae were fixed in 10% neutral buffer formalin for 48 hours and stored in 70% ethanol at 4 degrees Celsius.

### Dynamic Histomorphometry

A subset of humeri were embedded in polymethyl methacrylate (PMMA) and transversely sectioned under constant water irrigation via a low-speed precision saw at approximately 200 microns (0.0079 in) thick (IsoMet Low Speed Precision Cutter, Buehler, Lake Bluff, IL). Three transverse slices were taken per sample at the humeral diaphysis cutting from distal to proximal, fixed to slides with gorilla glue, sanded, and imaged at 20x magnification on a Zeiss LSM 880 laser scanning microscope with Airyscan (Carl Zeiss Microscopy, Thornwood, NY). Images were analyzed for standard dynamic histomorphometry metrics *(28)* including mineralized surface per bone surface (MS/BS), mineral apposition rate (MAR), bone formation rate (BFR/BS), total bone cross-section area (B.Ar), total mineralized area (Md.Ar), and total marrow area (Ma.Ar) using FIJI (built on ImageJ version 1.51n) and Photoshop (version CC 2020, Adobe Systems Inc., San Jose, CA). *Sample numbers were as follows: 7 postganglionic (6 for periosteal metabolism rates), 6 preganglionic, 5 sham*. Disarticulation could not be analyzed due to the unorganized woven bone that was laid down on the endosteal and periosteal surfaces, leaving no clear surface lines.

### Growth Plate Activity

The second subset of humeri was soaked in 30% sucrose solution for 24 hours and snap frozen in 2-methylbutance and stored at -80 degrees Celsius. Bones were cryosectioned longitudinally from medial to lateral using the Kawamoto method *(29)*. Briefly, two samples per bone were sectioned at -20 degrees Celsius by rolling cryofilm onto the sample before taking a 10 micron thick section. The cryofilm and sample were placed face up onto slides and adhered via UV adhesion glue.

After humeri samples were fixed on slides, images at 2.5x magnification (DMi8M, Leica Microsystems, Wetzlar, Germany) were taken surrounding the growth plate. Images were analyzed similar to previous dynamic histomorphometry performed. Bone growth rate was measured via FIJI (built on ImageJ version 1.51n) by calculating the distance between the top edge of calcein and alizarin labels and dividing it by the amount of time between injections (*7 days*). *Sample numbers were as follows: 7 postganglionic, 7 preganglionic, 6 disarticulation, and 7 sham*.

### Fibrosis

Snap frozen muscles were sectioned via cryotome (Cryotome FSE Cryostat, Thermo Scientific, Halethorpe, MD) at -20 degrees Celsius and 10 microns thick. Muscles included the biceps long head, biceps short head, upper subscapularis, and lower subscapularis, to identify altered muscle attached to the scapula and more accurately assess where fibrosis occurs in the biceps and subscapularis. Three tissue slices per muscle were taken from the belly and stained using a Masson’s trichrome staining kit (American MasterTech, Lodi, CA) for identification of collagen I. After imaging at 20x magnification (EVOS® FL Cell Imaging System, Thermo Scientific, Halethorpe, MD), analyses were performed in MATLAB (MATLAB®, The MathWorks, Inc., Natick, MA) using a custom processing protocol to calculate the percentage of collagen in the tissue section. Averages across the three sections were taken for each muscle.

Muscles used were half of the overall study. Due to some muscles being too small to section and some breakage, not all were analyzed. Sample numbers for affected limbs were as follows. *Postganglionic: 6 biceps short head, 8 biceps long head, 9 upper subscapularis, and 11 lower subscapularis. Preganglionic: 5 biceps short head, 10 biceps long head, 7 upper subscapularis, and 8 lower subscapularis. Disarticulation: 0 biceps short head, 0 biceps long head, 5 upper subscapularis, and 5 lower subscapularis. Sham: 7 biceps short head, 7 biceps long head, 7 upper subscapularis, and 7 lower subscapularis*. Sample numbers for unaffected limbs were as follows. *Postganglionic: 10 biceps short head, 10 biceps long head, 11 upper subscapularis, and 11 lower subscapularis. Preganglionic: 9 biceps short head, 10 biceps long head, 9 upper subscapularis, and 7 lower subscapularis. Disarticulation: 5 biceps short head, 5 biceps long head, 5 upper subscapularis, and 5 lower subscapularis. Sham: 4 biceps short head, 7 biceps long head, 7 upper subscapularis, and 6 lower subscapularis*.

### FGF-2 Quantification

ELISA assays were performed to quantify the amount of FGF-2 present in the biceps long head and upper subscapularis muscles. Due to limits on the number of samples we could use, the biceps short head and lower subscapularis were left out. Snap frozen muscles were homogenized in 2mL 1X PBS per 0.1g of muscle. Solutions were then frozen and thawed three times to lyse cells, centrifuged at 10 rpm for 10 min, and the supernatant taken for analyses. Assays were performed using an FGF-2 kit (LSBio, Seattle, WA) following the instruction manual. Sample numbers were as follows: *3 biceps long head and 3 upper subscapularis in postganglionic, preganglionic, and sham; 2 biceps long head and 2 upper subscapularis in disarticulation*.

### Statistical Analyses

First, normality of data was examined via D’Agostino-Pearson tests. Group comparisons between sham, postganglionic, preganglionic, and disarticulation were analyzed using either two-way ANOVA with Tukey’s post hoc tests or Kruskal-Wallis with Dunn posthoc, depending on normality of data. These analyses were run for each dynamic histomorphometry metric on both the endosteal and periosteal surfaces, growth plate activities, collagen content in each muscle, and FGF-2 quantifications in each muscle. Limb comparisons were then made between the affected and unaffected limbs within each group. Limbs were compared using paired parametric t-tests with Welch’s correction for unequal variances, or a Wilcoxon matched-pairs signed rank test depending on normality of data. All analyses were performed in Prism 6 (GraphPad Software, Inc., La Jolla, CA) with alpha = 0.05. Trends were defined as p < 0.10.

## Results

### Dynamic Histomorphometry

#### Limb Comparisons

The postganglionic group showed a trend towards limb differences in mineralization rates on the endosteal surface while the preganglionic group showed limb differences on the periosteal surface (Figure 1A, Table 1). On the endosteal surface, postganglionic tended to have lower mineralizing surface to bone surface (p=0.054, -16%), mineral apposition rate (p=0.064, -31%), and bone formation rate (p=0.078, -36%) on the affected side compared to the unaffected side. Preganglionic showed no differences on the endosteal surface. On the periosteal surface, preganglionic showed reduced mineral apposition rate (-28%) and bone formation rate (-34%), and tended to show reduced mineralizing surface to bone surface (p=0.094, -13%) on the affected side. However, postganglionic showed no differences on the periosteal surface.

**Figure 1.**
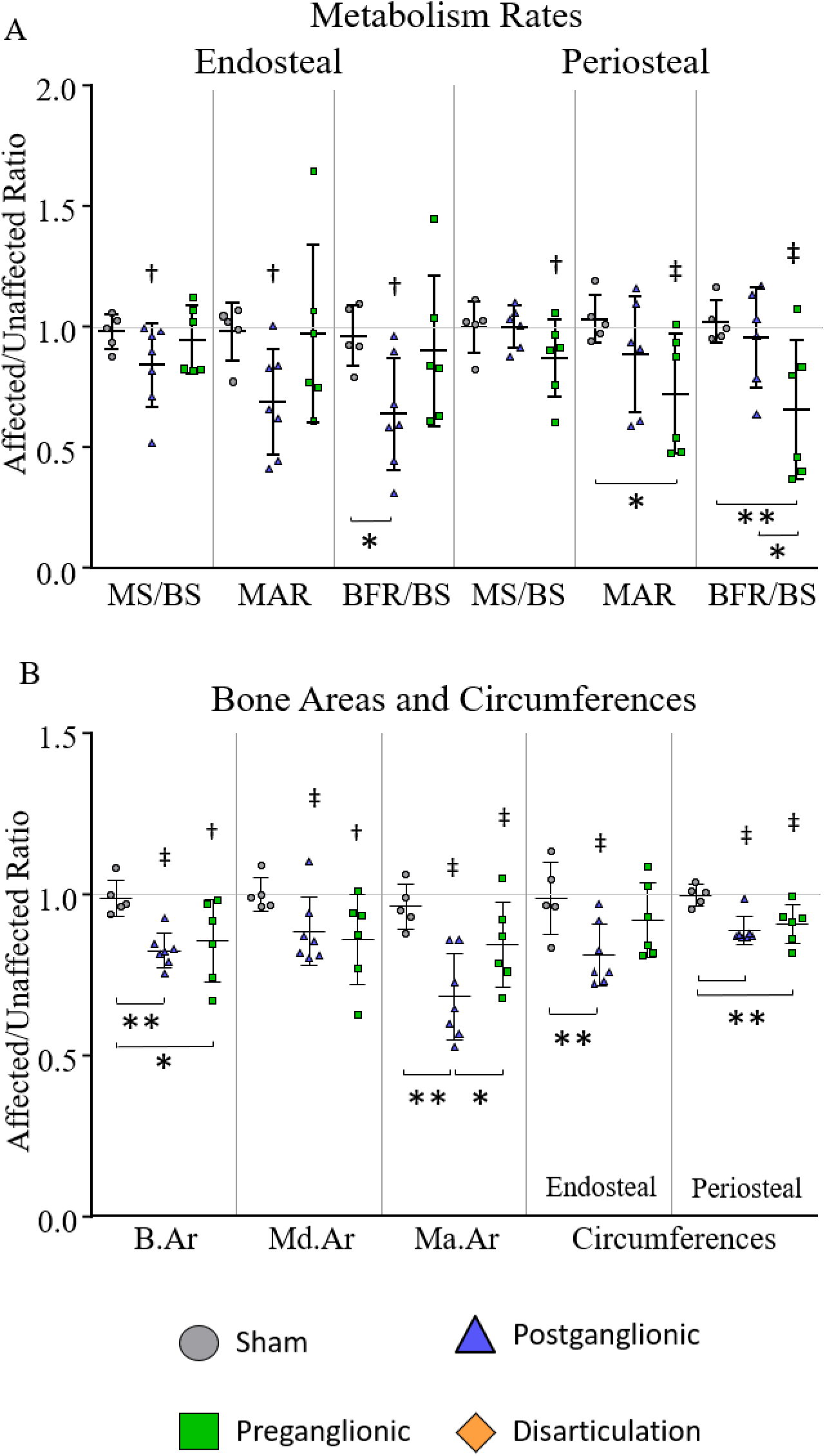
Bone metabolism rates (A) and cross-sectional areas (B) in the humeral mid-diaphysis following postganglionic and preganglionic BPBI. Postganglionic was affected more on the endosteal surface while preganglionic was affected more on the periosteal surface. ^‡^p < 0.05, ^†^p < 0.10 affected vs. unaffected limb; **p < 0.05, *p < 0.10 group differences.

**Table 1.**
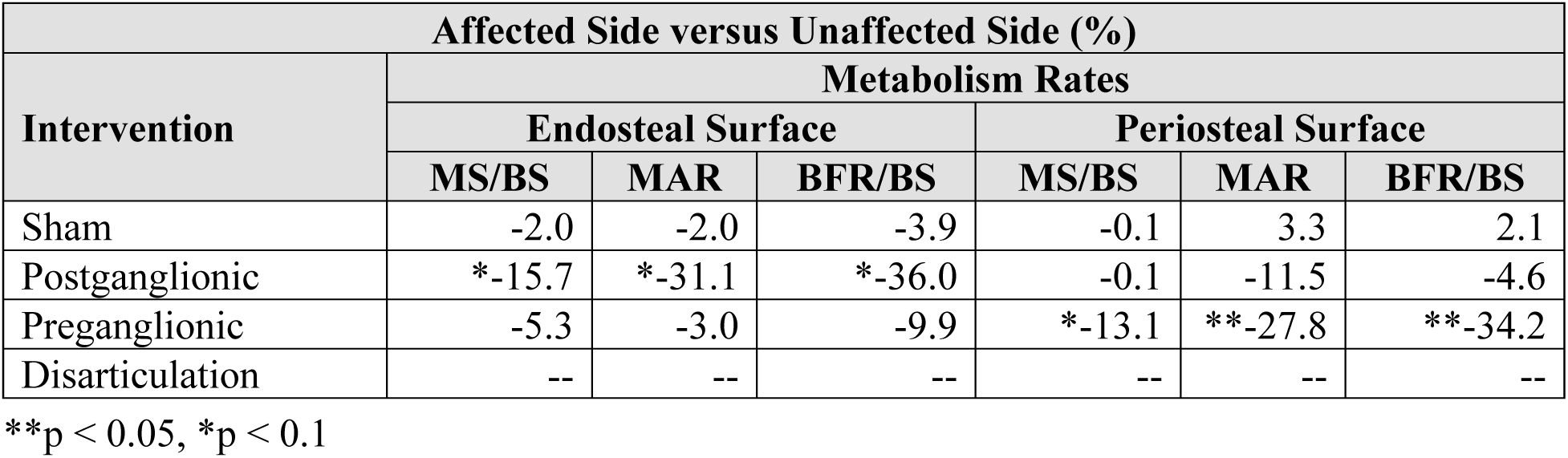
Percent differences in bone metabolism rates between affected and unaffected limbs in the humeral mid-diaphysis.

With regard to cross-sectional areas and surface circumferences, both postganglionic and preganglionic showed significant differences in the affected limb compared to the unaffected limb (Figure 1B, Table 2). Postganglionic had a 17% decrease in bone area, 11% decrease in mineralized area, 32% decrease in marrow area, 19% decrease in endosteal circumference, and 11% decrease in periosteal circumference. Preganglionic had 15% smaller marrow area, 9% smaller periosteal circumference, and tended to have a 14% smaller bone area (p=0.069) and mineralized area (p=0.096).

**Table 2.**
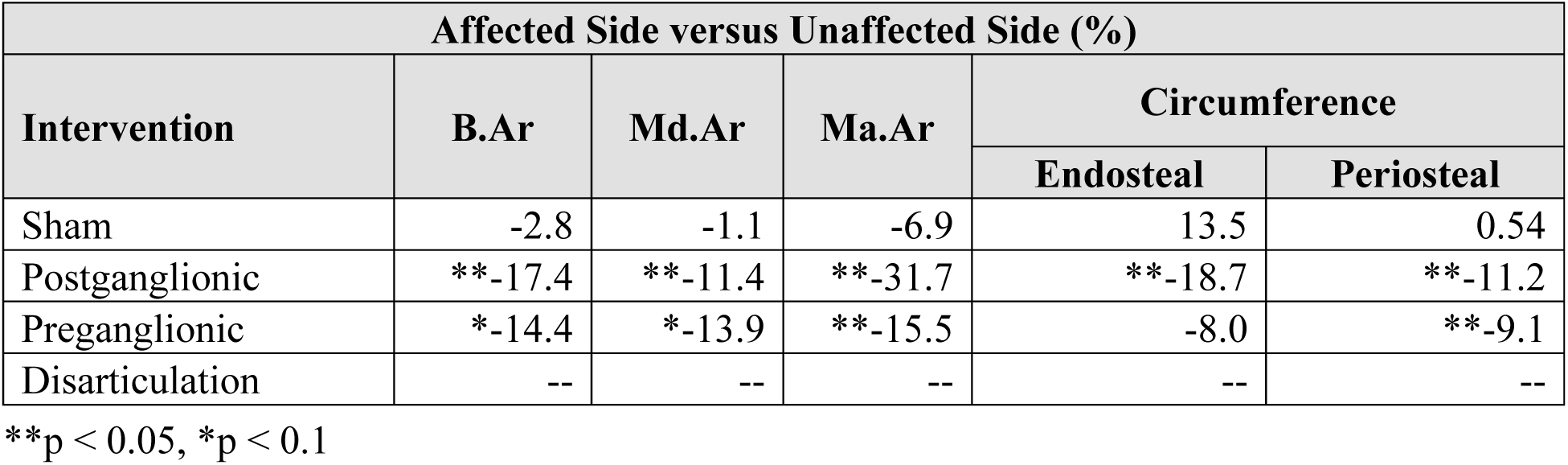
Percent differences in bone cross-sectional areas and circumferences between affected and unaffected limbs in the humeral mid-diaphysis.

#### Group Comparisons

The differences seen between groups were less than the limb differences within groups (Figure 1). On the endosteal surface, postganglionic injury tended to show 28% lower BFR/BS (p=0.090) compared to sham, but no significant differences were seen. On the periosteal surface, preganglionic showed a significant decrease in BFR/BS by 25% compared to sham. Preganglionic injury also tended to portray 17% lower MAR (p=0.071) compared to sham and 31% lower BFR/BS (p=0.078) compared to postganglionic.

Although preganglionic resulted in greater changes in mineralization rates, postganglionic resulted in more cross-sectional area and circumference changes. Postganglionic injury produced 17% and 12% smaller B.Ar and Ma.Ar respectively and 18% and 11% smaller endosteal and periosteal circumferences respectively compared to sham. Compared to preganglionic, postganglionic tended to produce a 24% smaller Ma.Ar (p=0.069). Preganglionic had a 9% smaller periosteal circumference and tended to have a 14% lower B.Ar (p=0.050) compared to sham.

### Growth Plate Activity

#### Limb Comparisons

Significant differences were seen between the affected and unaffected humeri (Figure 2, Table 3) in the preganglionic (-6%)and disarticulation (-6%) groups. The postganglionic group was not different enough to be significant.

**Figure 2.**
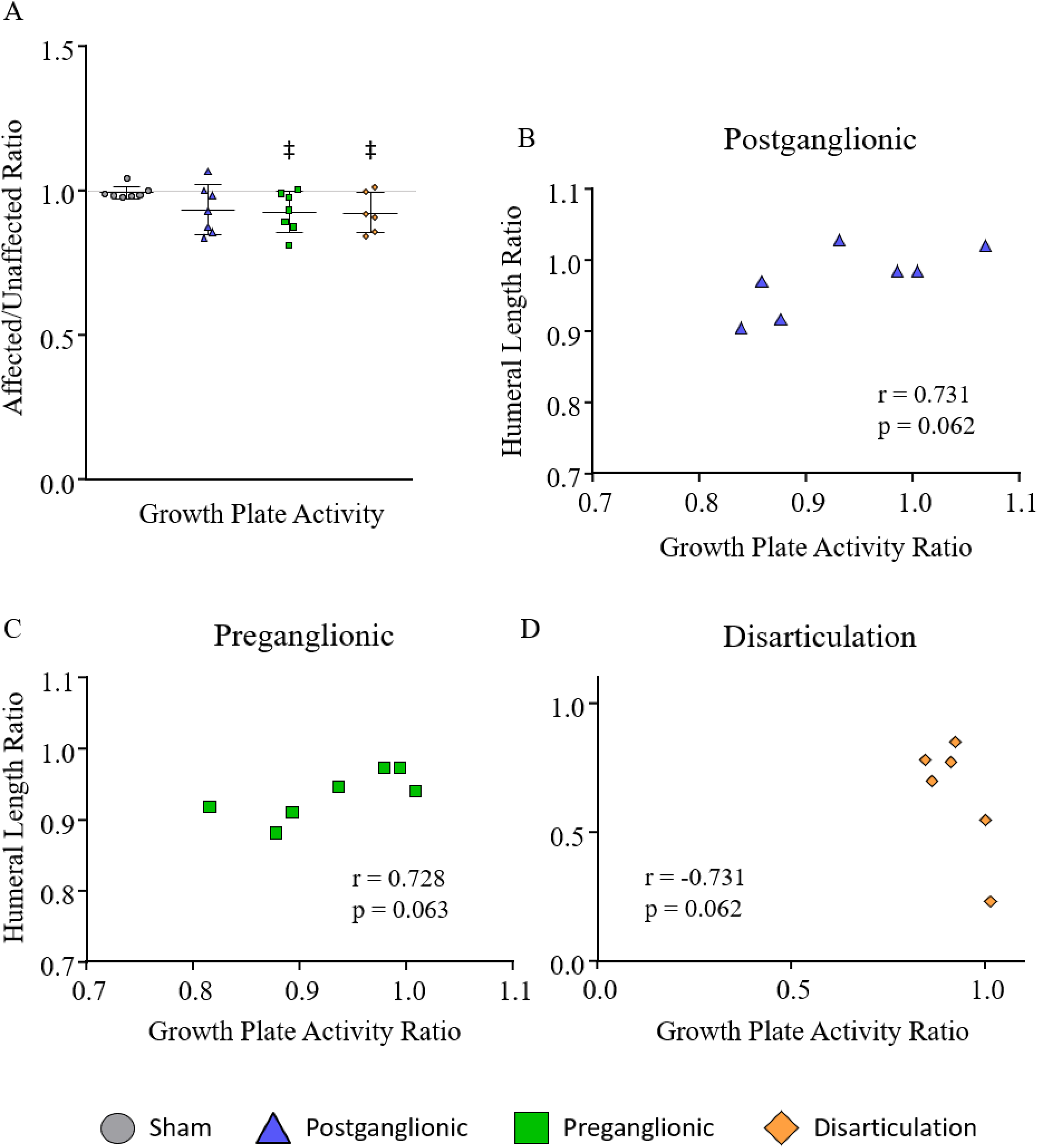
A) Activity of the proximal humeral growth plate. Preganglionic and disarticulation showed lower activity on the affected side. Correlations between humeral growth plate activity and humeral length were positive in the B) postganglionic and C) preganglionic groups and negative in the D) disarticulation group. ^‡^p < 0.05, ^†^p < 0.10 affected vs. unaffected limb; **p < 0.05, *p < 0.10 group differences.

**Table 3.** Percent differences in the rate of proximal humeral growth plate activity between the affected and unaffected sides.

| Affected Side versus Unaffected Side (%) |  |
| --- | --- |
| Intervention | Growth Plate Activity |
| Sham | 1.2 |
| Postganglionic | 1.0 |
| Preganglionic | *-6.2 |
| Disarticulation | *-5.8 |

\*\*p < 0.05, \*p < 0.1

#### Group Comparisons

No significant differences or trends were seen between groups (Figure 2). Although limb differences were seen with the preganglionic injury and disarticulation groups, the ratio of the limbs in each group was not significantly different from sham.

#### Correlations

While there were few differences seen in humeral growth plate activity, there were only trend towards significant correlations present within the postganglionic, preganglionic, and disarticulation groups when comparing humeral length and humeral growth plate activity (Figure 2). The postganglionic and preganglionic groups showed similar significantly positive correlations between the ratio of affected to unaffected humeral length and the ratio of affected to unaffected growth plate activity (p=0.062, r=0.731; p=0.063, r=0.728 respectively). As the difference in affected and unaffected humeral length increased, the difference in their respective growth plate activities also increased. However, the opposite was true for the disarticulation group. There was a trend towards a negative correlation (p=0.062, r=-0.731) in which the ratio of the affected to unaffected humeral length decreased as the ratio of the affected to unaffected growth plate activity increased.

### Fibrosis

Postganglionic and preganglionic injuries resulted in fibrosis in some of the muscles analyzed. Disarticulation showed no significant changes in collagen content (Figure 3).

**Figure 3.**
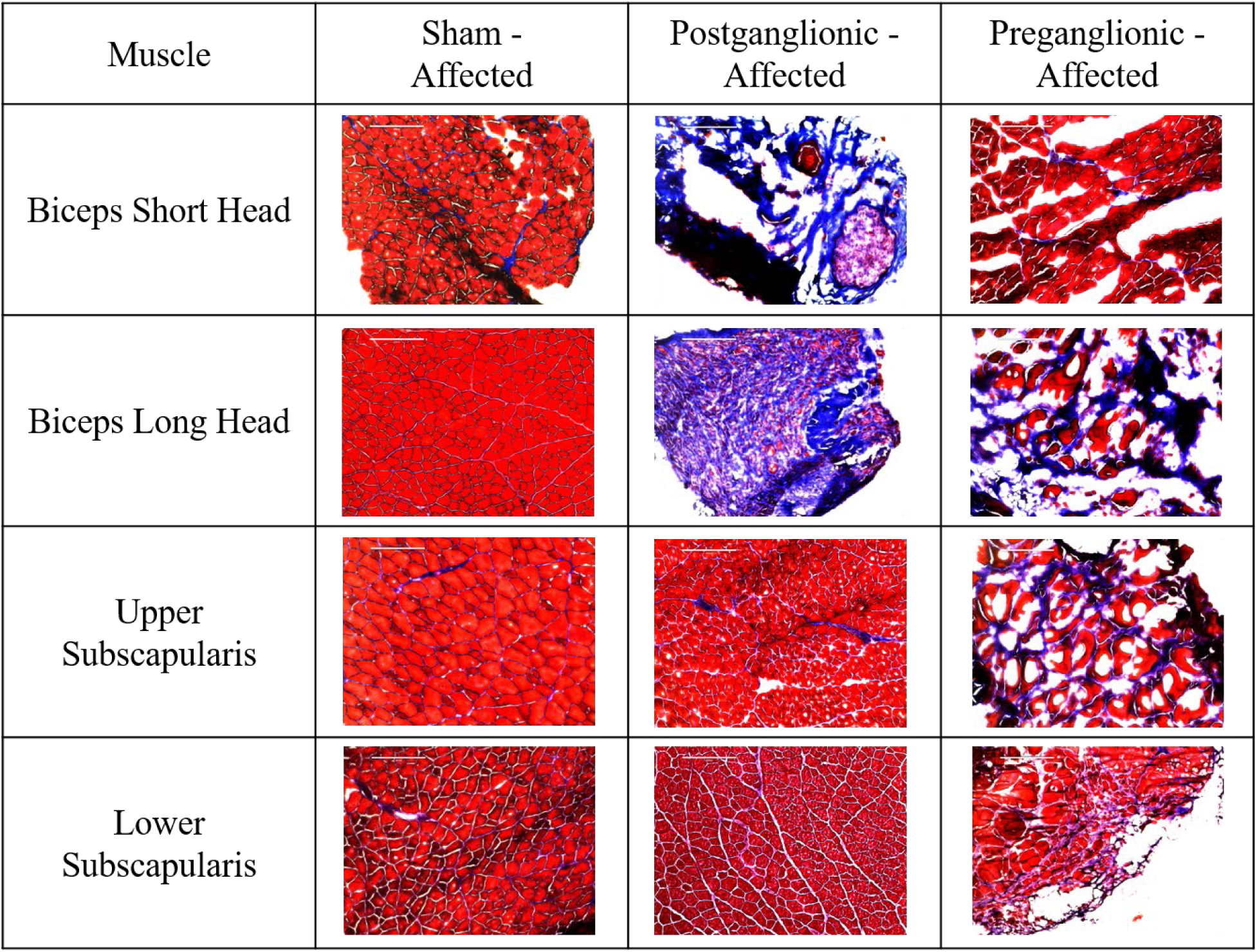
Chart with images representative of the collagen content in the affected limb following sham, postganglionic injury, and preganglionic injury.

#### Limb Comparisons

Few groups showed significant differences in fibrosis between affected and unaffected limbs (Figure 4, Table 4). Differences were seen in the biceps short head and biceps long head, but not in either the upper or lower subscapularis muscles. In the postganglionic group, the biceps short head of the affected limbs contained significantly higher collagen amounts (2.6x) than the unaffected limbs. In the preganglionic group, the biceps long head had significantly higher collagen content (3.1x) in the affected limbs compared to the unaffected.

**Figure 4.**
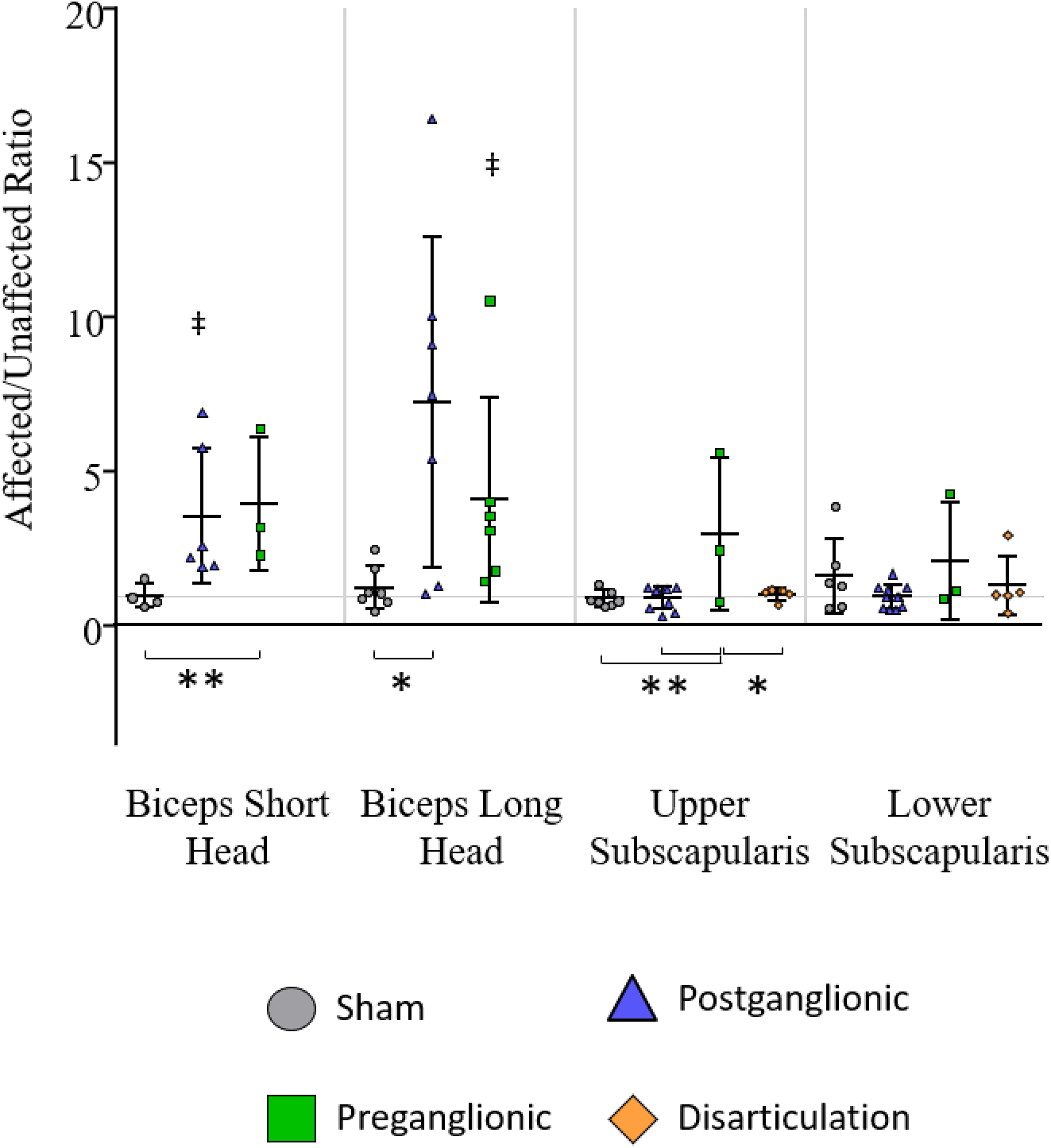
Collagen content in the biceps long head, biceps short head, upper subscapularis, and lower subscapularis. A) The ratio of collagen content in the affected limb to the unaffected limb. B) Collagen content on only the affected side of each group. C) Collagen content on only the unaffected side of each group. ^‡^p < 0.05, ^†^p < 0.10 affected vs. unaffected limb; **p < 0.05, *p < 0.10 group differences.

**Table 4.** Percent differences in muscle collagen content and FGF-2 quantity between affected and unaffected sides.

| Intervention | Affected Side versus Unaffected Side (%) |  |  |  |  |  |
| --- | --- | --- | --- | --- | --- | --- |
|  | Collagen Content |  |  |  | FGF-2 Quantification |  |
|  | Biceps |  | Subscapularis |  | Biceps | Subscapularis |
|  | Long Head | Short Head | Upper | Lower | Long Head | Upper |
| Sham | -3.5 | 23.3 | -11.7 | 62.2 | 0.7 | -1.1 |
| Postganglionic | 626.3 | **255.3 | -10.3 | -6.4 | 19.5 | -1.0 |
| Preganglionic | **307.2 | 296.5 | 195.8 | 109.7 | -4.4 | 4.9 |
| Disarticulation | -- | -- | 1.8 | 29.0 | -- | 2.5 |

\*\*p < 0.05, \*p < 0.1

#### Group Comparisons

Looking at the ratios of the affected limb to the unaffected limb, preganglionic had significantly more fibrosis in the biceps short head (2.5x) compared to sham and postganglionic showed higher fibrosis in the biceps long head (4.9x) compared to sham (Figure 4).

Similar differences in the upper and lower subscapularis were seen when comparing intervention groups. In the upper subscapularis, the ratio of affected to unaffected limbs was significantly increased in the preganglionic group compared to sham, postganglionic, and disarticulation (2.3x, 2.3x, and 1.9x respectively).

### FGF-2 Quantification

#### Limb Comparisons

No significant alterations were seen within any intervention groups for either muscle (Figure 5, Table 4). Although not significant, the average FGF-2 following postganglionic injury does appear to be larger in the affected biceps long head (19.5%). Additionally, both postganglionic and preganglionic groups appear to have increases in both the affected and unaffected limbs, explaining insignificant differences between the affected and unaffected limbs.

**Figure 5.**
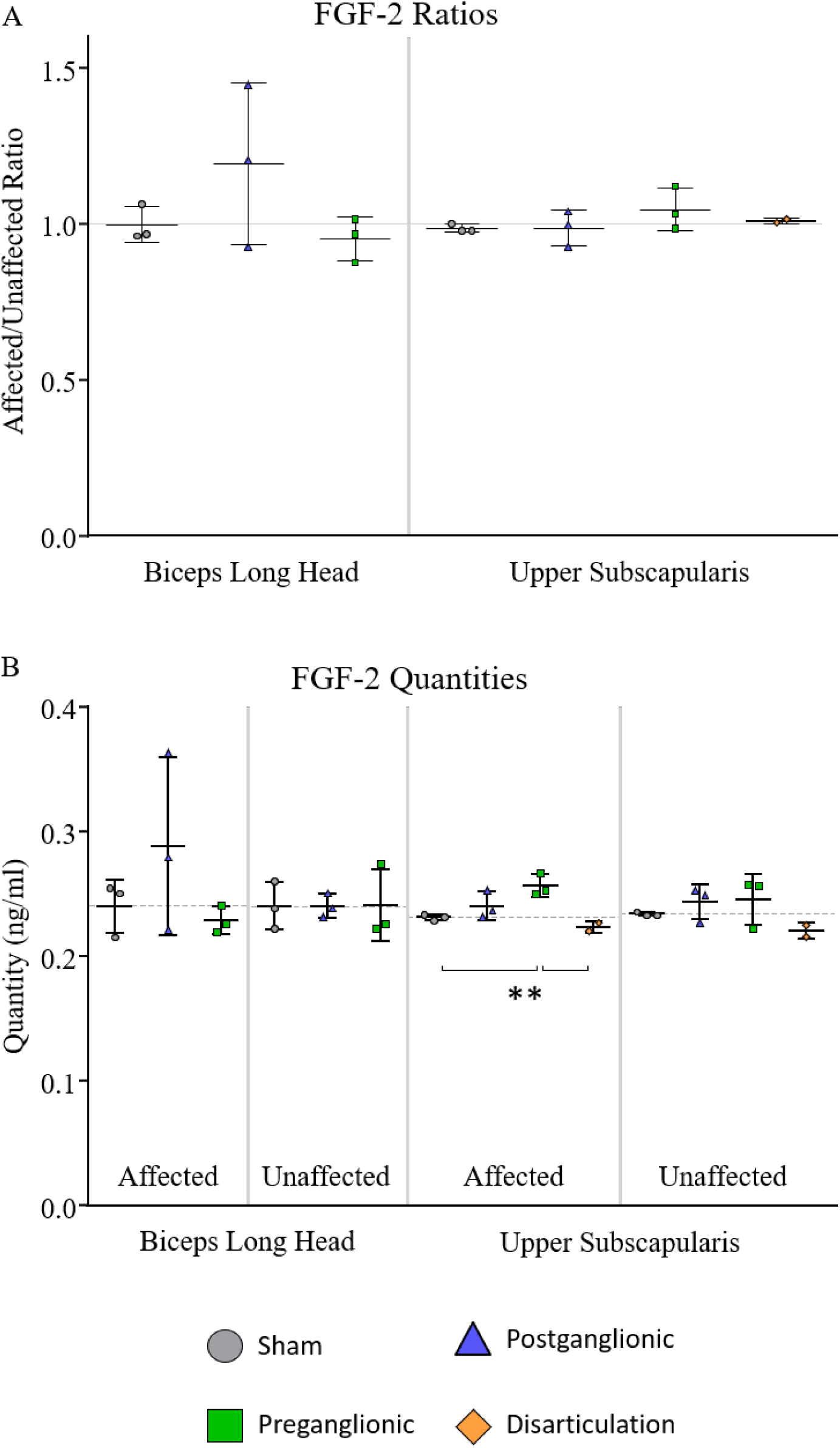
Quantification of FGF-2 in the biceps long head and upper subscapularis muscles. A) The ratio of FGF-2 quantity in the affected limb to the unaffected limb. B) FGF-2 content on the affected and unaffected sides within each group. ^‡^p < 0.05, ^†^p < 0.10 affected vs. unaffected limb; **p < 0.05, *p < 0.10 group differences.

#### Group Comparisons

Comparisons in the affected to unaffected ratios between groups showed no significant differences (Figure 5). However, the average of FGF-2 content appears higher after postganglionic in the biceps long head and after preganglionic in the upper subscapularis.

No significant differences were shown in the biceps long head between groups when comparing either the affected limbs or the unaffected limbs.

In the upper subscapularis, preganglionic had significantly higher FGF-2 content compared to sham and disarticulation (11.3% and 15.2% respectively). While no other comparisons were significant, there was a general appearance of systemically increased FGF-2 content in the upper subscapularis following both postganglionic and preganglionic injury. On the affected and unaffected sides for postganglionic there is an increase of 3.9% and 3.8% compared to sham respectively. On the affected and unaffected sides for preganglionic there is an increase of 11.3% and 4.7% compared to sham respectively.

Disarticulation aligned well with sham on all accounts.

## Discussion

The objective of this study was to determine bone metabolism, muscle composition, and muscle-bone crosstalk with regard to injury location and limb disuse following BPBI. Bone metabolism following the disuse model was unable to be examined due to the woven bone deposited throughout the humeral cortical shell rather than mature lamellar bone. Additionally, due to the procedure of amputation, the biceps long head and biceps short head could not be analyzed for collagen content or FGF-2 quantity in the disarticulation group. With regard to postganglionic and preganglionic injuries, slightly different patterns between the two emerged when looking at the results as a whole.

### Dynamic Histomorphometry

Bone growth and metabolism were altered differently during the seven-to-eight-week time points between the two neurectomy locations. Postganglionic injury resulted in more changes to overall diaphyseal regions and lower metabolism rates on the endosteal surface. Preganglionic injury resulted in less changes to overall diaphyseal regions and lower metabolism rates on the periosteal surface. This suggests postganglionic injury may be altered by both bone formation and bone resorption, which occur uncoupled on the endosteal surface during growth and development. The smaller cross-sectional areas of bone may also indicate reduced bone metabolism at an earlier time in life. Preganglionic shows altered bone formation of the outer surface during the seen-to-eight-week time, which is where osteoblasts lay most of the new bone during development.

The alterations on the endosteal surface following a postganglionic injury resulted in less area for the marrow cavity and a smaller endosteal circumference. The greater decrease of bone deposition on the periosteal surface following a preganglionic injury resulted in a smaller periosteal circumference and a trend towards lower overall cross-sectional area and lower mineralized area.

These results suggest preganglionic injury causes an overall reduction in bone growth. During growth and development, the periosteal surface is where the majority of the osteoblast activity occurs with regard to cortical bone. This explains the smaller humeral head in previous measures of morphology *(8, 12)*. The lower metabolism also implies diminished growth of trabecular bone, which aligns with results displayed earlier. While altered mechanical loading on the joint is also likely causing alterations in trabecular bone following both postganglionic and preganglionic injuries, there is the additional factor of bone metabolism.

### Growth Plate Activity

Results of the growth plate activity followed a similar qualitative pattern with the previous data collected on humeral length in the same cohort of rats. Postganglionic, preganglionic, and disarticulation had similar decreases of activity in the affected arm with preganglionic and disarticulation being significant. When correlating the growth plate activity and humeral length data, postganglionic and preganglionic showed a trend of moderate positive correlation between the two, suggesting the altered metabolism of the proximal growth plate following neurectomy causes changes in humeral length. In the disarticulation group, there was a trend of a negative correlation between the metrics. However, it is likely that the amputation performed for this group interfered with the humeral length data, making it unclear if the correlation is true. Additionally, the rat pups were very young with uncalcified bones, making it easy to amputate the distal growth plate with the rest of the arm. Either situation adds additional unknown factors into the correlation.

Overall, the growth plate activity was similar to the humeral length data. The postganglionic group had some decrease in growth plate activity, but not enough to be significant. With a larger sample size, the data may show a greater decrease in postganglionic and have a significant correlation between the two data metrics. In preganglionic and disarticulation groups, additional data may show significant changes rather than trends.

In the current data, the growth rates from the proximal humeral growth plate suggest reduced metabolism and growth in the preganglionic injury. It also suggests postganglionic deformity may in part, be due to reduced growth. This particular longitudinal shortening may be related to disuse of the affected limb.

### Fibrosis

Collagen content was not increased by different degrees in the biceps and subscapularis muscles. No changes were seen in the biceps short head, but in the biceps long head, both postganglionic and preganglionic injuries resulted in an increase in collagen content. In the subscapularis muscles, no changes were seen following postganglionic injury or disarticulation. However, preganglionic injury showed some increase in both the upper and lower subscapularis.. With regard to the disarticulation group, no increase in collagen content was seen in any muscles analyzed, suggesting limb disuse does not play a role in the development of fibrosis, regardless of injury location.

While collagen content was not increased as much in the biceps as the subscapularis, it was increased following both postganglionic and preganglionic injuries, making it consistent between injury locations. The subscapularis had greater alterations but only after a preganglionic injury. These results suggest limb loading is not responsible for fibrosis in the biceps or subscapularis muscles, and therefore other biochemical drivers are present following injury. It also suggests that although preganglionic injury preserves afferent innervation and does not present with contracture, more fibrosis is developed than after a postganglionic injury. Therefore, consistent with previous studies *(21)*, fibrosis cannot have a causal effect on contracture.

Roger Cornwall’s group also showed fibrosis in the biceps muscles that was similar between postganglionic and preganglionic injuries *(21)*. However, the subscapularis muscles (along with other related muscles) had not been examined. This study evaluated the subscapularis muscle because of its architectural changes and its involvement in altered mechanical loading of the joint. Clinical studies agree that the subscapularis has the most severe changes of all the shoulder muscles *(2, 30, 31)* and the results of this study align well as the subscapularis had more fibrosis than the biceps following a preganglionic injury. While the subscapularis could cause altered loading on the joining, the muscle composition changes themselves cannot be due to altered limb loading, suggesting deformity contributions from nerve injury.

### FGF-2 Quantification

The biceps long head did not show any significant differences in FGF-2 between or within intervention groups. In the upper subscapularis, no significant changes were seen in either postganglionic or disarticulation groups. However, preganglionic injury did result in a small increase of FGF-2, which was likely systemic rather than only increased on the affected side.

Seeing significant results from this data is less likely due to the low number of samples. Looking at the qualitative pattern of the data in isolation of statistical significance, it appears as though postganglionic and preganglionic injuries have a systemic increase in the upper subscapularis with preganglionic being worse. It also appears that there could be an increase FGF-2 of the affected biceps long head following a postganglionic injury. The disarticulation group was most similar to sham.

Currently obtained data for FGF-2 quantity indicates preganglionic injury has altered production of muscle cell-signaling while postganglionic and disarticulation do not. Therefore, the subscapularis may be more involved in injury than the biceps. Additionally, preganglionic injury may have a driver, other than disuse, which is not affecting postganglionic injury.

The addition of more data, however, may reveal trends of upregulated FGF-2 in the upper subscapularis for both postganglionic and preganglionic injury. This would suggest, again, that the subscapularis is more involved than the biceps and that a nerve injury itself is affecting both nerve injury locations, but more so in the preganglionic.

As seen with other peripheral nerve injuries, FGF-2 was upregulated (27, 32, 33) in the subscapularis following a preganglionic injury, despite the fibrosis present. This suggests fibrosis does not interfere with muscle-cell signaling with regard to FGF-2 generation in muscle. It also implies the preservation of afferent nerves contributes to the ability of the subscapularis to upregulate growth factors, similar to conclusions drawn from the preservation of ErbB signaling seen after a preganglionic injury *(5)*. Upregulation of FGF-2 after injury aids in muscle recovery and is therefore a positive consequence *(27, 32, 33)*. After a postganglionic injury, ErbB signaling was disrupted *(5)* and aligns with the results found here considering there was no significant increase in FGF-2 as is normally present after injury. This suggests that the ability of the muscle to generate an appropriate amount of cell-signaling protein, specifically FGF-2, is related to the preservation of the afferent nerves. However, if preserving afferent nerves also preserves FGF-2 cell-signaling ability, then a reduction in FGF-2 is not likely the cause of metabolism changes seen prior to osseous deformity in a preganglionic injury model.

### Limitations

Although this study furthered the knowledge of how bone is altered and potentially why it is altered, there were some limitations. In the animal models, the nerve is severed rather than stretched or torn like in humans. Additionally, rats are quadrupedal their entire lives whereas humans are quadrupedal for a short amount of time. However, even with these slight differences between the animal models and humans, it has been shown that the deformities following the rat models recapitulate the deformities seen in humans *(8)*. The second limitation in this study is the sample numbers, especially regarding the FGF-2 ELISAs. Not all the samples could be used for all parts of these analyses due to differing techniques and fixations of samples. However, there were still some significant trends that were revealed, giving insight to potential underlying mechanisms of osseous deformity. Lastly, the amputation performed on the disarticulation group damaged the biceps muscles crossing the elbow joint. Therefore, the biceps short head and biceps long head could not be analyzed for the disarticulation group.

## Conclusions

Overall, the results of this study suggest that osseous deformities after a preganglionic injury result from reduced bone growth by a combination of limb disuse and nerve injury. The osseous deformity after a postganglionic injury likely results more from disuse. Of the potential mechanisms causing deformity, muscle fibrosis and FGF-2 production in muscle are not likely. Fibrosis is present after each injury type but does not correspond to osseous deformities seen. Additionally, preganglionic injury portrays more fibrosis, suggesting the cause of fibrosis differs between the two interventions. Preganglionic injury likely has a greater increase because of the greater stunt in growth of muscles compared to postganglionic injury. FGF-2 is also upregulated after preganglionic injury, which may suggest better cell-signaling after a preganglionic injury compared to postganglionic. Additionally, disuse of the affected limb does not contribute to fibrosis or cell-signaling, suggesting separate underlying factors. One additional factor that should be investigated includes satellite cells. The extra-cellular matrix (ECM), where fibrosis occurs, and satellite cells affect one another. Altered quantity of satellite cells in the muscle may explain the fibrosis, upregulation of FGF-2, and previously seen architectural changes which affect bone growth.

## Acknowledgments

This work was supported by the National Institutes of Health [R21 HD088893] and the National Science Foundation [DGE-1746939].

